# Effect of tryptophan depletion on conditioned threat memory expression: role of intolerance of uncertainty

**DOI:** 10.1101/2020.04.26.062505

**Authors:** Jonathan W Kanen, Frederique E Arntz, Robyn Yellowlees, David M Christmas, Annabel Price, Annemieke M Apergis-Schoute, Barbara J Sahakian, Rudolf N Cardinal, Trevor W Robbins

## Abstract

**Background:** Responding emotionally to danger is critical for survival. Normal functioning also requires flexible alteration of emotional responses when a threat becomes safe. Aberrant threat and safety learning occurs in many psychiatric disorders including post-traumatic stress disorder (PTSD), obsessive-compulsive disorder (OCD), and schizophrenia, where emotional responses can persist pathologically. Whilst there is evidence that threat and safety learning can be modulated by the serotonin systems, there have been few studies in humans. We addressed a critical clinically relevant question: How does pharmacological lowering of serotonin affect the retention of conditioned threat memory?

**Methods:** Forty-seven healthy participants underwent threat conditioning on Day 1 followed by an extinction session. Emotional responding was assessed by the skin conductance response (SCR). On Day 2, we employed acute dietary tryptophan depletion to lower serotonin temporarily, in a double-blind placebo-controlled randomized between-groups design. We then tested for the return of conditioned threat memory spontaneous recovery). We also measured self-reported intolerance of uncertainty, known to modulate threat memory expression.

**Results:** The expression of emotional memory was attenuated in participants who had undergone tryptophan depletion. Individuals who were more intolerant of uncertainty showed even greater attenuation of emotion following depletion.

**Conclusions:** These results support the view that serotonin is involved in predicting aversive outcomes and refine our understanding of the role of serotonin in the persistence of emotional responsivity, with implications for individual differences in vulnerability to psychopathology.

## Introduction

The ability to respond emotionally to threats in the environment is critical for an organism to optimise behavior and navigate the world. Once a threat is no longer present, it is crucial to adapt emotional responses flexibly to reflect the safe environment, for normal functioning in daily life to continue. Dysfunction of threat and safety learning lies at the core of post-traumatic stress disorder (PTSD; Milad et al., 2009) and other anxiety disorders (Kim et al., 2011; Marin et al., 2017), and is also a feature of obsessive-compulsive disorder (OCD; Apergis-Schoute et al., 2017; McLaughlin et al., 2015; Milad et al., 2013) and schizophrenia (Holt et al., 2012). PTSD is unique amongst these in that exposure to a traumatic event is a defining feature, and it is characterised by subsequent pathological physiological reactions to cues reminiscent of the event (APA, 2013). Elucidating the factors that contribute to the persistence of such emotional reactions is essential in order to develop new treatments. Here we tested the influence of the neuromodulator serotonin (5-hydroxytryptamine; 5-HT) on the retention of emotional memory, with a widely used laboratory model of PTSD and related disorders (Graham and Milad, 2011).

Pavlovian threat conditioning paradigms involve pairing a previously neutral stimulus with an aversive outcome, such as a mild electric shock. Individuals learn that the cue signals threat, and an anticipatory sympathetic nervous system arousal response occurs. This manifests as measurable perspiration and is known as the skin conductance response (SCR). After an individual has learned that a cue signals threat, the stimulus can later be repeatedly presented without the aversive consequence (extinction learning) - a model of exposure therapy in the clinic by which a new memory of safety should be formed. These two memories - of threat and safety - then compete for expression upon re-encountering a conditioned stimulus. Threat memories are well known to persist regardless of extinction training, and the re-emergence of the emotional memory after the passage of time is known as spontaneous recovery (Bouton, 2002). Emotional memories for threats can also resurface following re-exposure to an adverse event, known as reinstatement (Bouton, 2002). Understanding what contributes to spontaneous recovery and reinstatement is of great clinical interest, because of its implications for conditions such as PTSD (Graham and Milad, 2011; Milad and Quirk, 2012). One factor that has recently come to light is self-reported intolerance of uncertainty (IUS; Carleton et al., 2007): individuals highly intolerant of uncertainty demonstrated greater spontaneous recovery (Dunsmoor et al. 2015).

Serotonin, meanwhile, is widely implicated in aversive learning (Cools et al., 2008), and several studies have begun to explore the role of serotonin in threat and safety learning, and aversive memory (Bauer, 2015). Most experiments, however, have been carried out in rodents (Bauer, 2015). The dearth of human studies at the nexus of threat memory and serotonin function is particularly surprising, given that first-line pharmacological treatments of disorders in which threat conditioning processes are impaired - e.g. PTSD, OCD, and other anxiety disorders - modulate serotonin (Stahl, 2013). No one, to our knowledge, has manipulated serotonin experimentally to examine its influence on spontaneous recovery in humans.

Acute tryptophan depletion (ATD) is a commonly used method for studying serotonin function in which tryptophan, the amino acid biosynthetic precursor to serotonin, is temporarily removed from the diet in the presence of other amino acids. This results in decreased serotonin synthesis (Bel & Artigas, 1996; Biggio et al., 1974; Crockett et al., 2012; Nishizawa et al. 1997; Young 2013). ATD can modulate threat conditioning in humans: Hindi Attar et al. (2012) showed that ATD attenuated aversive conditioning, whilst Hensman et al. (1991) demonstrated that the 5-HT2A and −2C antagonist ritanserin also abolished conditioned threat responses, both assessed by SCR.

Serotonin can also have influences beyond initial learning. Two weeks’ administration of the serotonin reuptake inhibitor (SRI) escitalopram in humans did not impact the acquisition of threat memory but facilitated extinction (Bui et al., 2012). Karpova et al. (2011) demonstrated in mice that chronic administration of the SRI fluoxetine, when paired with extinction training, diminished spontaneous recovery and reinstatement. Hartley et al. (2012) used a behavioural genetics approach in humans and found a relationship between spontaneous recovery and normal variation in the serotonin transporter polyadenylation polymorphism (STPP) - whereas variation in the more widely studied 5-HTTLPR (5-HT-transporter-linked polymorphic region) had no effect. Additionally, neither polymorphism impacted the acquisition or extinction of threat memory. It should be noted, however, that studies of single nucleotide polymorphisms are vulnerable to false positives (Border et al. 2019), which in turn reinforces the importance of pharmacological approaches. The present study of healthy human volunteers investigated two key questions: how does pharmacologically lowering serotonin affect the return of conditioned threat memory? Does self-reported intolerance of uncertainty influence emotion? We employed ATD to investigate this clinically relevant question and predicted that lowering serotonin would modulate the expression of a previously formed threat memory.

## Methods and Materials

### Participants

Forty-seven healthy participants (mean age 25; 29 males) who met criterion for Pavlovian conditioning as assessed by the skin conductance response (SCR) were included in this experiment (see Supplemental Information). Participants were medically healthy and screened to be free from any psychiatric disorders. Individuals who reported, during screening, having a first-degree relative (parent or sibling) with a psychiatric disorder were also excluded. Participants reported not taking any regular medication (aside from contraceptive pills), had never taken psychiatric or neurological medications, and did not have any neurological conditions. See Supplemental Information for further criteria. Participants gave informed consent before the start of the study and were paid for their participation.

### General Procedure

The protocol was approved by the Cambridge Central Research Ethics Committee (UK National Health Service Ethics reference 16/EE/0101). Participants attended sessions on two consecutive days, which took place at the National Institute for Health Research / Wellcome Trust Clinical Research Facility at Addenbrooke’s Hospital in Cambridge, England. Day 1 comprised a short afternoon session with no pharmacological manipulation. Participants completed the first part of the Pavlovian task, baseline questionnaires, and two other unrelated non-emotional computer tasks not reported here. To assess mood and other feelings including alertness, a 16-item visual analogue scale (VAS) was administered at the beginning of day 1, and at the beginning, middle, and end of the main testing session on day 2. On Day 2 participants arrived in the morning having fasted for at least 9 hours. Participants then completed the VAS, gave a blood sample, and ingested either the placebo or ATD drink. In the afternoon participants completed the Pavlovian task, along with multiple other tasks that will be reported elsewhere.

### Acute Tryptophan Depletion

Tryptophan is the necessary amino acid precursor to synthesize brain serotonin. Acute tryptophan depletion (ATD) is a widely used dietary manipulation, which results in a rapid decrease in the synthesis of serotonin (Bel & Artigas, 1996; Biggio et al., 1974; Crockett et al., 2012; Nishizawa et al. 1997). Participants were randomly assigned to receive either ATD or a placebo condition, in a double-blind, between-groups design. The depletion group received a drink that contained a balance of all the essential amino acids except for tryptophan. The placebo group received the same drink except it included tryptophan (Worbe et al., 2014). Blood plasma samples were collected to verify tryptophan depletion.

### Pavlovian Conditioning Task

The task design was adapted from Hartley et al. (2012) and Milad et al. (2009) and is depicted in Figure 1. The key dependent measure was SCR, and participants were subjected to the threat of mild transcutaneous electrical stimulation (shock). On day 1, participants underwent a calibration procedure to determine a shock level that was “uncomfortable, but not painful” to them. They then underwent a Pavlovian threat conditioning procedure: two coloured square images (2 conditioned stimuli [CSs], denoted CS+E and CS+N) were presented for 4 seconds, on 16 trials each, and paired with receipt of shock (unconditioned stimulus; US) on 37.5% (6 of 16) of the presentations, while another coloured square image (CS−) was presented for 10 trials and was never paired with the US. For the CS+ stimuli, the US occurred 3800 milliseconds after stimulus onset, and lasted 200 milliseconds, co-terminating with the image. There was an inter-trial interval that averaged 10 seconds. After a few trials of this procedure it is normal for participants to show an anticipatory arousal response (reflected by mild perspiration of the fingers and measured by the SCR) upon viewing the CS+s, relative to the CS−. SCR measurement allowed verification that Pavlovian threat learning had occurred.

**Figure 1.**
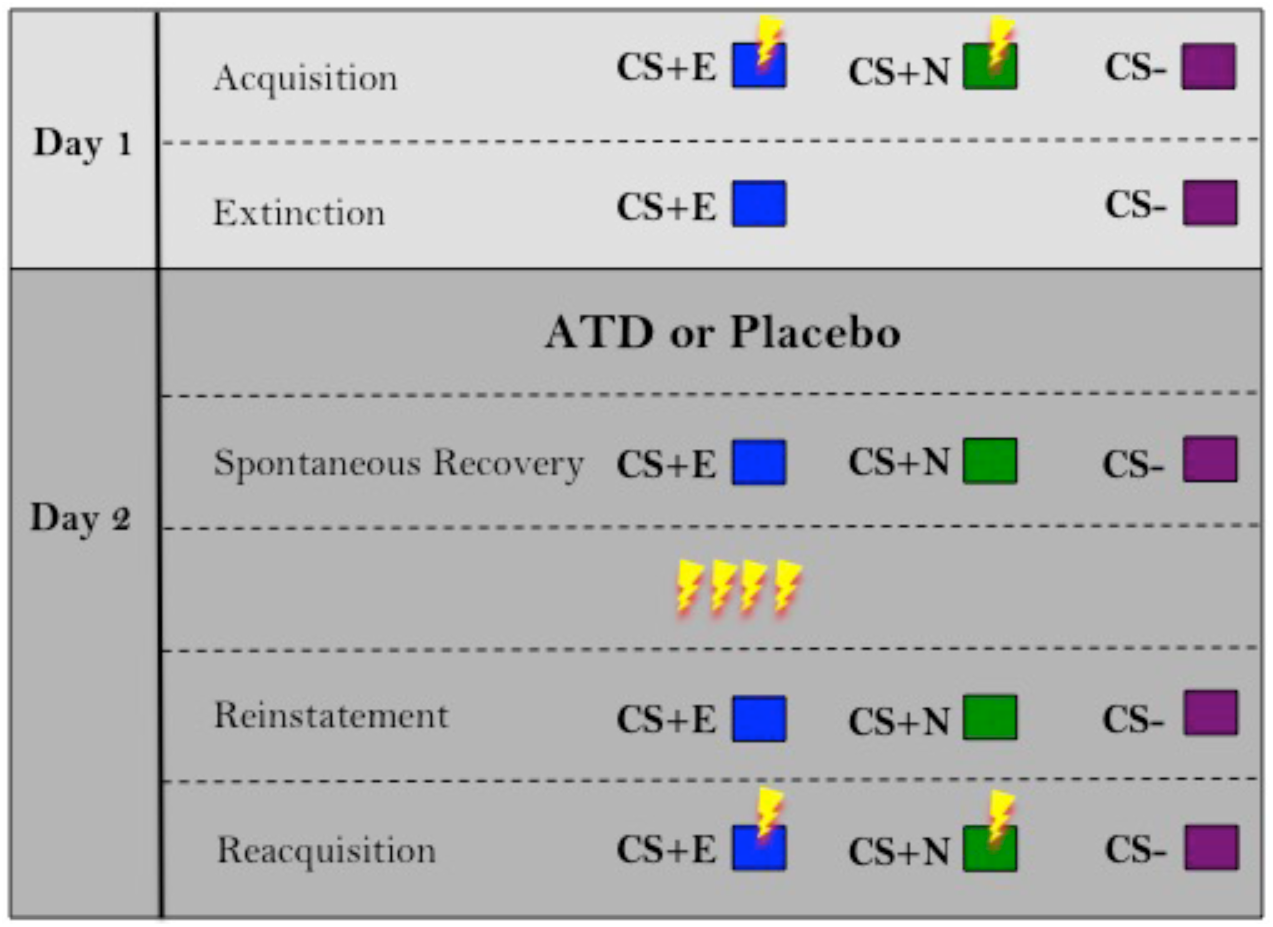
Task schematic. Each row represents a different phase of the experiment. Lightning bolts represent shock. CS+E is the conditioned stimulus that was presented during the extinction phase. CS+N is the CS+ that was not presented during the extinction phase. The CS− was never paired with shock. ATD = acute tryptophan depletion.

Participants then underwent standard extinction training, in which they were repeatedly presented with the CS+E (E for extinguished) and the CS−, both without the US. The other CS+ (the CS+N, N for not extinguished) was not presented during the extinction session.

Participants then returned one day later to receive ATD or placebo and were subsequently tested for the return of conditioned threat memory, using SCR across three phases. In the first phase, participants were first re-exposed to all three CSs (CS+E, CS+N, CS−), again without the US, to assess spontaneous recovery to the CS+E - the return of threat memory after the passage of time (Bouton, 2002). At this stage, the CS+N is a comparator against which spontaneous recovery of the CS+E can be measured. If ATD modulates expression of the original threat memory, it would be expected to alter responses to the CS+N. If it specifically affects the expression of the extinction memory (spontaneous recovery), it would be expected to alter responses to the CS+E but not the CS+N. In phase two, four USs were administered that were not associated with any of the images. Participants were subsequently re-exposed participants to all three images (reinstatement). The last phase of the experiment was a reacquisition procedure, where the CS+E and CS+N were once again paired with the US on 37.5% of trials (the CS− was also presented, without shock, as before). Greater reacquisition can be reflective of a stronger threat memory (Bouton, 2002). Importantly, the context remained the same across both days.

## Results

### Analysis of plasma tryptophan

Twenty-five participants underwent tryptophan depletion, whilst the remaining 22 received placebo. Plasma samples were analysed using high performance liquid chromatography (HPLC). The ratio of tryptophan to large neutral amino acids (TRP:LNAAs; valine, methionine, isoleucine, leucine, tyrosine, and phenylalanine) was calculated, as this is thought to be most reflective of the extent of brain 5-HT depletion (Fernstrom, 1979). A t-test was performed on the change in the TRP:LNAA ratio between samples taken at baseline and approximately 4.5 hours following administration of the mixture. Plasma levels were unavailable for two participants: one due to a staff processing error, and one due to unsuccessful venepuncture. A robust depletion of tryptophan was achieved (t_(43)_ = − 15.317, p = 5.05×10^−19^), displayed in Figure 2.

**Figure 2.**
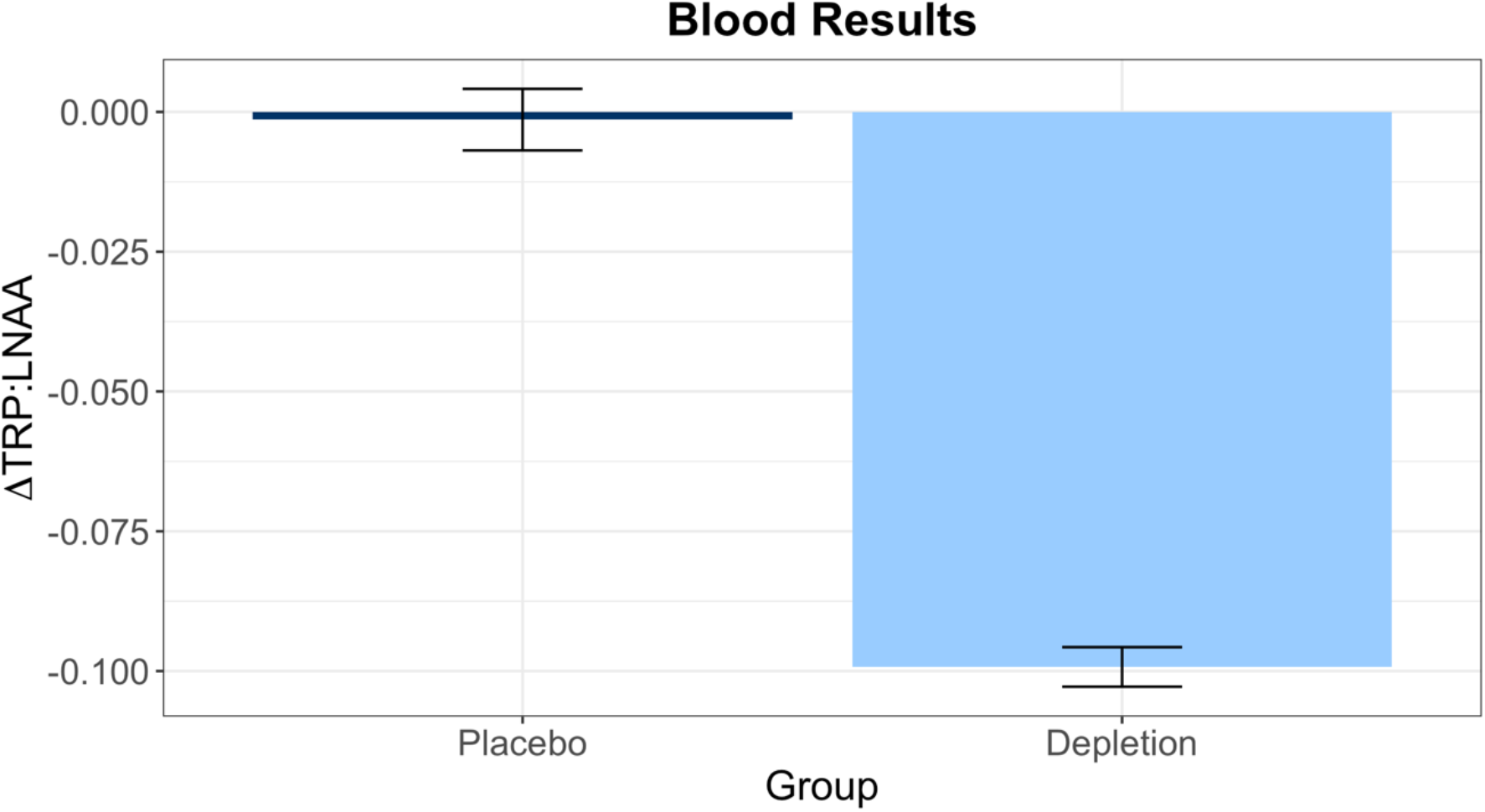
Robust tryptophan depletion was achieved, verified via plasma samples. ΔTRP:LNAA = change in the ratio between tryptophan and all large neutral amino acids before and after depletion. More negative bar indicates greater depletion of tryptophan. Error bars indicate 1 standard error.

### Self-report measures

Rating data was collected from 40 participants (n = 21 depletion) on how happy or sad they were feeling prior to the task, after depletion had taken effect, and these mood ratings did not differ from those participants who received placebo (t_(38)_ = −1.227, p = .228). At baseline, participants completed a number of other questionnaires which assessed depressive symptoms, anxiety, intolerance of uncertainty, among other measures, which are summarised in Table 1.

**Table 1.** Group characteristics

|  | **mean (SD) placebo** | **mean (SD) depletion** |
| --- | --- | --- |
| Age | 24.23 (5.88) | 25.80 (6.24) |
| Years edu | 17.18 (2.38) | 17.28 (2.42) |
| BDI-II | 5.23 (4.38) | 4.04 (3.92) |
| STAI | 37.36 (7.27) | 35.52 (5.96) |
| IUS | 55.73 (13.69) | 49.80 (13.44) |

NART = National Adult Reading Test (Nelson, 1982), a proxy for intelligence; BDI-II = Beck Depression Inventory (Beck et al., 1996); STAI = Spielberger Trait Anxiety Inventory (Spielberger et al., 1983); IUS = Intolerance of Uncertainty Scale (Carleton et al., 2007); edu = education; SD = standard deviation.

### Acquisition of Pavlovian threat conditioning before tryptophan depletion

Pavlovian threat conditioning, which was conducted without any serotonergic manipulation, was successfully achieved on day 1. Importantly there was no difference in conditioning on day 1 between those who later (on day 2) received placebo versus depletion, shown in Figure 3. A repeated measures ANOVA with group assignment (future placebo versus future ATD) and stimulus (CS+E, CS+N, CS−; all trials) as factors yielded a main effect of stimulus (F_(1, 61)_ = 22.031, p = 2×10^−6^, η_p_^2^ = .383), no main effect of group assignment (F_(1,45)_ = 0.378, p = .542, η_p_^2^ = .008) and no group assignment-by-stimulus interaction (F_(1, 61)_ = 0.658, p = .466, η_p_^2^ = .014). Follow up paired t-tests confirmed the CS+E (t_(46)_ = −5.315, p = 3×10^−6^) and CS+N (t_(46)_ = −4.632, p = 3×10^−5^) mean SCR values were each significantly greater than the SCR to the CS−.

**Figure 3.**
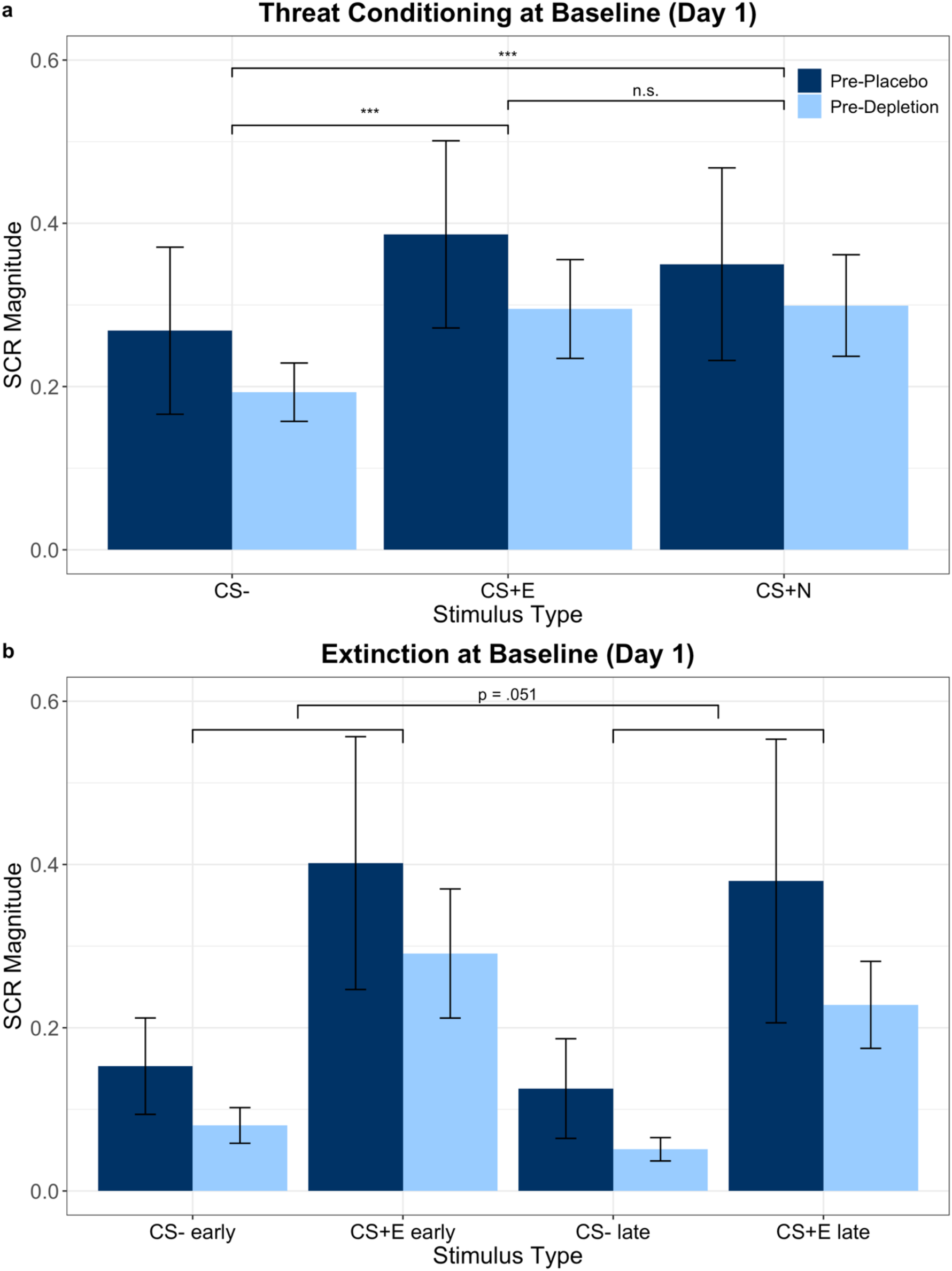
Skin conductance responses (SCR) in the **(a)** initial threat conditioning phase (acquisition) on Day 1, conducted before serotonergic challenge. There were no differences between the future placebo and future ATD groups and both groups showed significant threat conditioning to both CS+s compared to the CS−, as predicted. This equivalent baseline conditioning on Day 1 enabled testing the effects of ATD on its retention on day 2. Brackets denote follow-up t-tests contrasting stimuli within group, after observing a main effect of stimulus; *** indicates significance at p < .001; n.s. = not significant; error bars indicate 1 standard error. **(b)** SCR in the extinction phase on Day 1. Smaller brackets refer to the beginning and end of extinction, and the larger bracket denotes a marginally significant reduction in SCR in late compared to early extinction; error bars indicate 1 standard error.

### Extinction before tryptophan depletion

Extinction, which was also conducted before any serotonergic manipulation, was assessed by comparing the mean of the first two and mean of the last two trials, for each of the two stimuli presented during extinction. As expected, there was no difference during extinction on Day 1 between those who later (on Day 2) received placebo versus depletion (F_(1,45)_ = 0.932, p = .340, η_p_^2^ = .020), nor was there an interaction between group and phase (early vs. late trials; F_(1,45)_ = 0.364, p = .549, η_p_^2^ = .008). We expected that by the end of extinction, CS+E responses would no longer be significantly different from CS− responses, however full extinction was not achieved: there was no phase (early vs. late) by stimulus (CS+E vs. CS−) interaction (F_(1,45)_ = 0.187, p = .668, η_p_^2^ = .004) and the main effect of stimulus persisted (F_(1,45)_ = 11.217, p = .002, η_p_^2^ = .2). There was, however, a marginally significant effect of phase (F_(1,45)_ = 4.004, p = .051, η_p_^2^ = .082), showing lower responding, irrespective of stimulus, in the late trials.

### Spontaneous recovery following tryptophan depletion

Spontaneous recovery was the critical test in this experiment to address the main hypothesis that serotonin modulates the expression of previously formed Pavlovian threat memories. Indeed, ATD modulated emotional responses during the spontaneous recovery phase (Figure 4). To assess whether ATD affected the expression of the conditioned memories from Day 1, SCR during the first half of the spontaneous recovery phase on Day 2 was examined. Repeated measures ANCOVA was conducted with serotonin status (placebo versus ATD) and stimulus (CS+E, CS+N, CS−) as factors, controlling for the strength of initial conditioning, and intolerance of uncertainty. SCR during acquisition was used as a covariate because we were interested in assessing the influence of the pharmacological manipulation on memory expression irrespective of how the strength of the initial memory affected expression a day later. Scores from the intolerance of uncertainty scale (IUS) were used as an additional covariate because this trait can affect threat memory expression (Dunsmoor et al., 2015). The repeated measures ANCOVA yielded a significant main effect of serotonin status (F_(1,43)_ = 8.818, p = .005, η_p_^2^ = .170), showing that emotional responses were significantly attenuated under ATD. There was also a main effect of stimulus (F_(2,73)_ = 3.594, p = .040, η_p_^2^ = .077). Follow up paired t-tests revealed responses to the CS+E and CS+N collapsed across serotonergic status were each significantly greater than responses to the CS- (t_(46)_ = −4.549, p = 3.9×10^−5^ and t_(46)_ = −5.089, p = 7×10^−6^, respectively), which demonstrated return of threat memory expression occurred irrespective of the serotonergic manipulation. Responses to the CS+E and CS+N did not differ from one another (t_(46)_ = −0.312, p = .756) which is likely because there was not complete extinction of the CS+E on day 1. There was no serotonin × stimulus interaction (F_(1,72)_ = 1.795, p = .179, η_p_^2^ = .040), indicating the effect of ATD was not specific to any of the three stimuli. Conditioning to both CS+s from Day 1 was retained on Day 2 in both the placebo and ATD groups; however, overall emotional responsivity was diminished by ATD, irrespective of stimulus. Threat memory expression, as measured by SCR in the spontaneous recovery phase, was not abolished by ATD but was attenuated.

**Figure 4.**
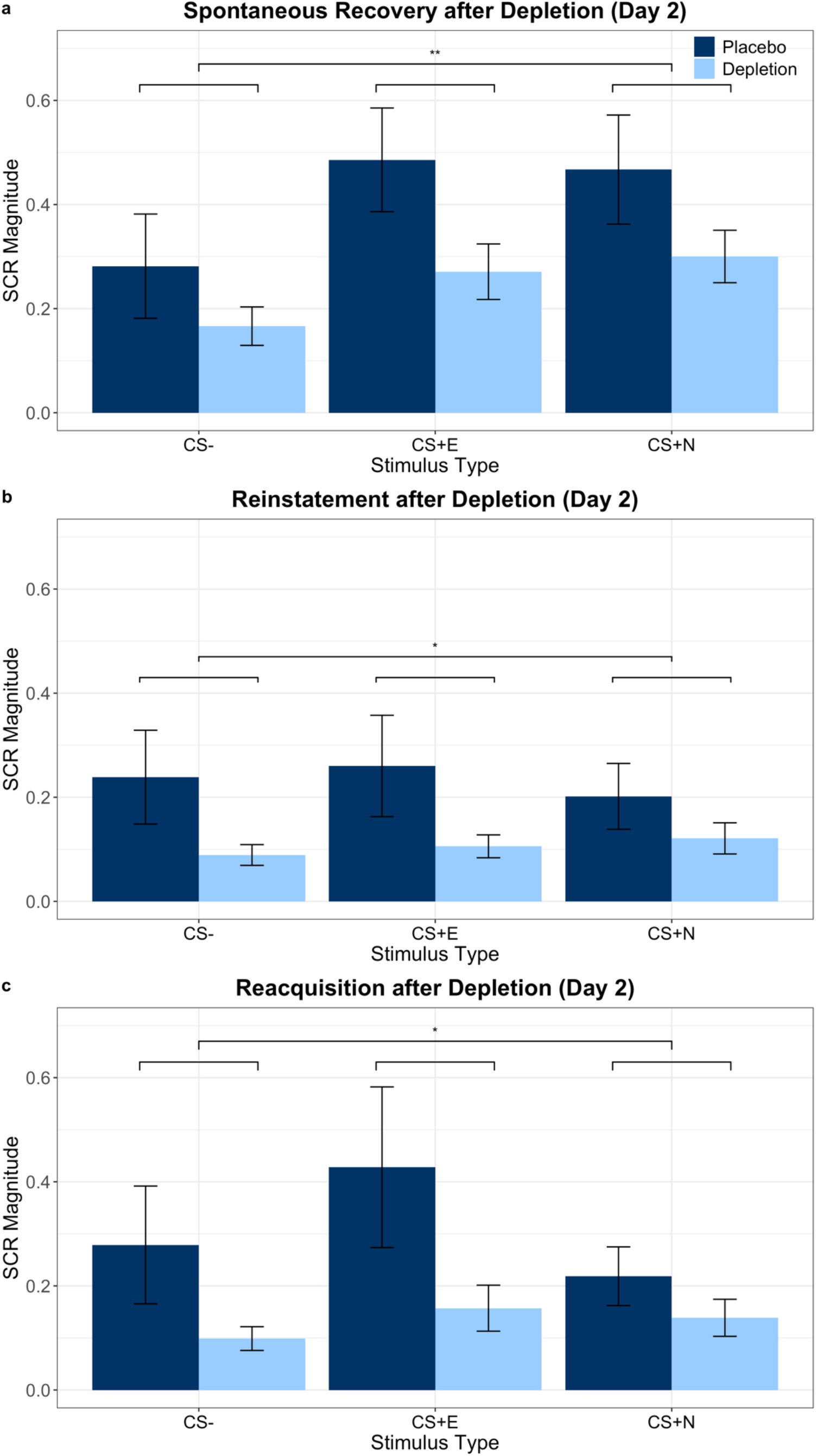
Tryptophan depletion reduced SCR expression. Skin conductance responses (SCR) are displayed from the **(a)** spontaneous recovery, **(b)** reinstatement, and **(c)** reacquisition phases. Large brackets denote a main effect of stimulus; ** indicates significance at p < .01; *indicates a significance at p < .05. Error bars indicate 1 standard error. Raw data are displayed, not adjusted values after controlling for intolerance of uncertainty or strength of initial conditioning on Day 1.

### Role of intolerance of uncertainty in the effects of ATD on spontaneous recovery

Next we examined how trait intolerance of uncertainty contributed to the results in the spontaneous recovery phase. Correlation analyses were performed between IUS and SCR to each of the stimuli, controlling for strength of Day 1 conditioning. These analyses revealed significant correlations in the depletion group only: when under depletion, individuals who were more intolerant of uncertainty showed significantly diminished emotional expression to the CS+E (r_(25)_ = −.554, p = .004) and CS+N (r_(25)_ = −.453, p = .023), as well as the CS- (r_(25)_ = −.418, p = .038). Under placebo this relationship with IUS was not present for any of the three stimuli: CS+E (r_(25)_ = −.135, p = .549), CS+N (r_(25)_ = - .249, p = .264), and CS- (r_(25)_ = −.109, p = .629). Critically, these results remained significant after being subjected to the Benjamini-Hochberg procedure for six comparisons at q = .15, as used in Skandali et al. (2018). Next, an interaction term between serotonin and IUS was incorporated into the general linear model used in the initial analysis of spontaneous recovery, in order to examine whether trait intolerance of uncertainty and serotonin status interacted to modulate SCR to specific stimuli. ANCOVA with serotonin and IUS as a between-subjects interaction term, controlling for main effects and strength of initial conditioning, and stimulus (CS+E, CS+N, CS−) as within-subjects factors did not show a two-way interaction between serotonin and IUS (F_(1,42)_ = .056, p = .815, η_p_^2^ = .001) or a three-way interaction between serotonin, IUS, and stimulus (F_(2,69)_ = 1.379, p = .257, η_p_^2^ = .032). Whilst there was no interaction effect between ATD and IUS, the correlation results suggest that ATD modulated the relationship between IUS and SCR to the conditioned stimuli.

### Relationship between spontaneous recovery and extent of tryptophan depletion

The extent of tryptophan depletion significantly correlated with the attenuation of threat responding, but not safety memory expression during the spontaneous recovery phase, which is displayed in Figure 5. Critically, this substantiated the relationship between depletion and conditioned threat memory expression during spontaneous recovery. Using a partial correlation to control for strength of acquisition and intolerance of uncertainty, there was a significant relationship between degree of tryptophan depletion and the extent to which the threat memory returned. Extent of depletion correlated with SCR to the CS+E and CS+N, and not the CS−, indicating the effect of tryptophan depletion did not generalise to safety memory expression: threat memory responses were more attenuated, the greater the depletion (CS+E, r_(41)_ = .409, p = .006; CS+N, r_(41)_ = .390, p = .01; CS−, r_(41)_ = .107, p = .495), These results additionally survived the Benjamini-Hochberg procedure for three comparisons, at q = .15 (Skandali et al. 2018). There was no interaction between stimulus (CS+E, CS+N, CS−) and plasma results on SCR, however, as assessed by repeated measures ANCOVA with plasma values, intolerance of uncertainty, and strength of initial conditioning as predictors (F_(2,66)_ = .890, p = .397, η_p_^2^ = .022).

**Figure 5.**
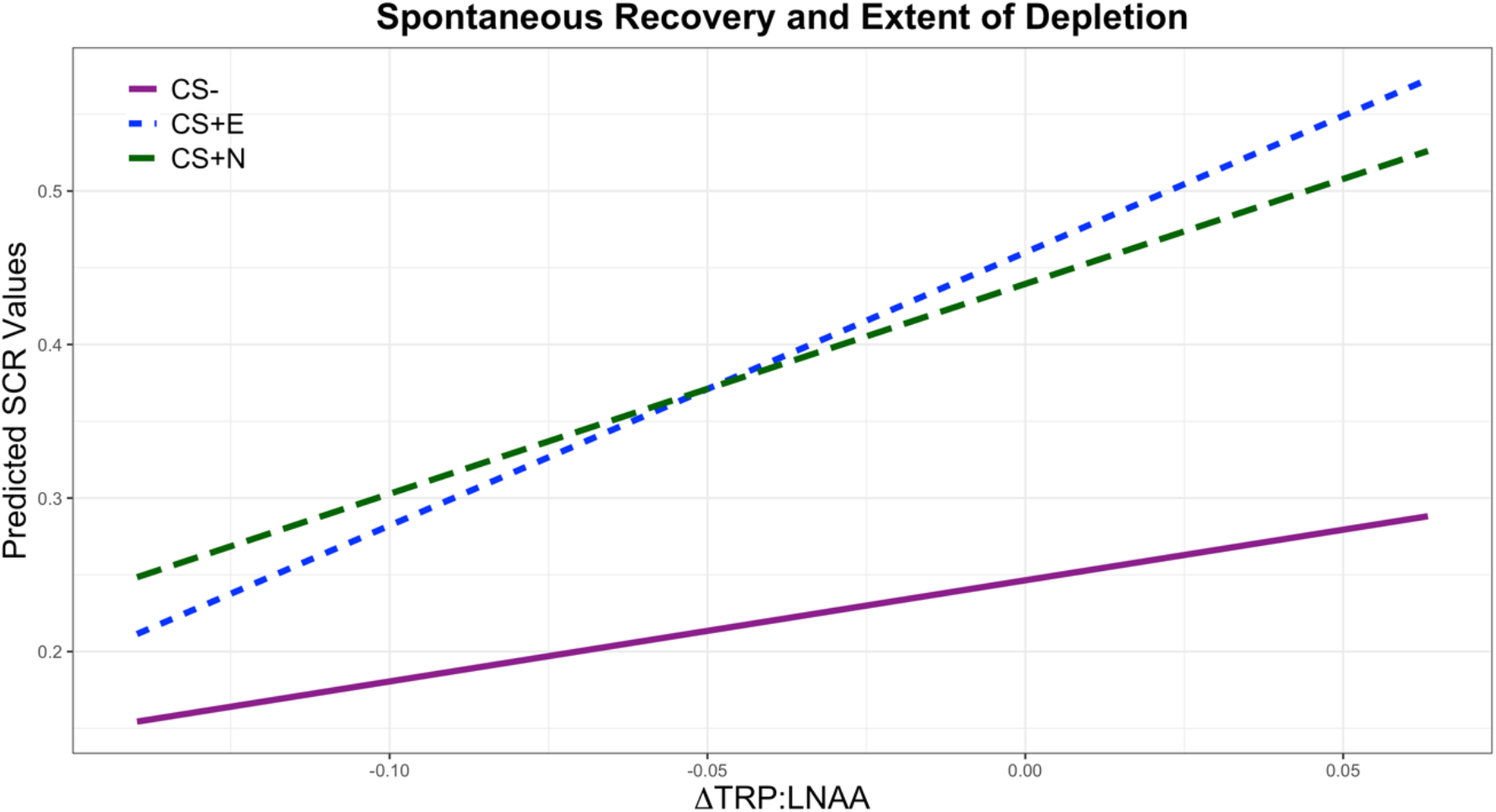
Predicted values for spontaneous recovery plotted against the extent of depletion assessed via plasma samples. Lower x-axis values indicate greater depletion. Predicted values were generated based on a univariate ANCOVA where the dependent variable was SCR to each stimulus during the spontaneous recovery phase, and the predictors were intolerance of uncertainty, strength of conditioning on Day 1, and the change (Δ) in the ratio of tryptophan (TRP) to large neutral amino acids (LNAA).

### Effects of tryptophan depletion on unconditioned responses

Responses to the unconditioned stimulus were not modulated by ATD. The unconditioned response (UR) to the shock (US) was analysed to evaluate whether ATD reduced emotional responsiveness in general, or whether the attenuation was specific to cue-evoked emotion. To do this, SCR to the four US presentations during the reinstatement procedure (where shocks were presented as discrete events, unpaired to any CS) were isolated and averaged. Two participants were not included in this analysis and those that follow: one due to an equipment failure, one due to experimenter error. Responses to the reinstatement USs were not affected by ATD (t_(43)_ = −0.897, p = .375), which was also the case when controlling for the magnitude of URs on Day 1 (at baseline) and intolerance of uncertainty (F_(1,41)_ = 1.240, p = .272, η_p_^2^ = .029). URs on Day 2, moreover, were not correlated with the extent of depletion, controlling for magnitude of URs on Day 1 and intolerance of uncertainty (r_(39)_ = .268, p = .091).

### Effects of tryptophan depletion on reinstatement

Tryptophan depletion modulated emotional responses in reinstatement. The reinstatement phase was analysed in the same way as the spontaneous recovery phase, examining the first half of trials, and controlling for strength of acquisition and intolerance of uncertainty. Repeated measures ANCOVA revealed a significant effect of serotonin status (F_(1,41)_ = 5.578, p = .023, η_p_^2^ = .120), again with lower emotional responses under ATD. By this stage in the experiment, however, there was no longer a main effect of stimulus (F_(1, 51)_ = 0.068, p = .849, η_p_^2^ = .002); there was no differential response to the CS+s relative to the CS−, and thus no reinstatement of the threat memory. Effects in these paradigms are often short-lived (Gershman & Hartley 2015). There was also no serotonin × stimulus interaction (F_(1, 51)_ = 1.390, p = .251, η_p_^2^ = .033).

### Relationship between reinstatement and extent of tryptophan depletion

To explore further explore the relationship between ATD and responses in reinstatement, a partial correlation analysis was conducted. Controlling for strength of acquisition and intolerance of uncertainty, there was a significant correlation between depletion and CS+E and CS- responses, but not CS+N responses (CS+E, r_(39)_ = .399, p = .01; CS+N, r_(39)_ = .232, p = .144; CS-, r_(39)_ = .452, p = .003). This is likely reflective of the fact that at this stage in the experiment there was no longer differential conditioning to the CS+s versus CS−.

### Effects of tryptophan depletion on reacquisition

For the final phase of the experiment, the same analysis used in the spontaneous recovery and reinstatement phases was repeated, which examined the first half of trials. Repeated measures ANCOVA showed a non-significant effect of serotonin status (F_(1,41)_ = 3.442, p = .071, η_p_^2^ = .077). There was no main effect of stimulus (F_(1, 60)_ = 1.915, p = .166, η_p_^2^ = .045) nor a serotonin × stimulus interaction (F_(1, 60)_ = 0.114, p = .832, η_p_^2^ = .003). Therefore, there was no evidence of re-conditioning in either the placebo or depletion groups. In both groups, however, the CS+E was numerically greater than the CS−, while the CS+N and CS- were numerically similar. Because effects in these paradigms are often short-lived (Gershman & Hartley 2015), and participants’ SCR tended to habituate later in the experiment, the same analysis was repeated on the first two trials of the re-acquisition phase which then showed a main effect of serotonin status: responses to the CSs were attenuated overall by ATD (F_(1,41)_ = 6.946, p = .012, η_p_^2^ = .145). There was no main effect of stimulus (F_(2,82)_ = 1.610, p = .206, η_p_^2^ = .038), nor a serotonin × stimulus interaction (F_(2,82)_ = 2.421, p = .095, η_p_^2^ = .056), again providing no evidence of re-conditioning in either group. In both groups, however, the CS+E was again numerically greater than the CS−, while the CS+N and CS- were numerically similar.

### Relationship between reacquisition and extent of tryptophan depletion

A partial correlation analysis was performed, as in the prior phases. Focusing on the first half of trials in the reacquisition phase, there were significant correlations between the extent of depletion and responses to each of the three stimuli (CS+E, r_(39)_ = .338, p = .031; CS+N, r_(39)_ = .352, p = .024; CS-, r_(39)_ = .336, p = .032). This partial correlation analysis was repeated, isolating the first two trials, as before. There was a significant correlation between depletion and the SCR response to the CS+E and CS−, but not the CS+N (CS+E, r_(39)_ = .472, p = .002; CS+N, r_(39)_ = .227, p = .154; CS−, r_(39)_ = .367, p = .018). This is again likely reflective of the fact that at this stage in the experiment there was no longer differential conditioning to the CS+s versus CS−.

### Relationship with trait anxiety

Intolerance of uncertainty was highly correlated with trait anxiety (r_(47)_ = .529, p = 1.32 × 10^−4^). An additional ANCOVA was therefore conducted, with serotonin status (placebo, depletion) as between-subjects factors, stimulus (CS+E, CS+N, CS−) as within-subjects factors, with trait anxiety, intolerance of uncertainty, and strength of conditioning on Day 1 as covariates. Whilst including trait anxiety as an additional covariate in an ANCOVA assessing spontaneous recovery (the primary phase of interest in this study) reproduced the key main effect of serotonin status (F_(1,42)_ = 8.473, p = .006, η_p_^2^ = .168), trait anxiety was not a significant predictor over and above intolerance of uncertainty (F_(1,42)_ = .742, p = .394, η_p_^2^ = .017). Intolerance of uncertainty, on the other hand, was a significant predictor over and above trait anxiety (F_(1,42)_ = 4.423, p = .041, η_p_^2^ = .095). When incorporating STAI into the model, moreover, there was no longer a simple main effect of stimulus (F_(2,71)_ = 2.634, p = .088, η_p_^2^ = .059).

### Summary of results

There was successful baseline (Day 1) Pavlovian threat conditioning, which did not differ between groups. There were also no baseline differences in extinction. The key result was that ATD attenuated the expression of previously acquired emotion in the spontaneous recovery phase. Accounting for self-reported intolerance of uncertainty (IUS), furthermore, contributed to the prediction of how ATD modulated SCR on Day 2. Differential conditioning, meanwhile, was not abolished by ATD. Whilst the reduction in SCR during the spontaneous recovery phase by ATD was not specific to any of the three stimuli at the group level, the greater the extent of tryptophan depletion, the more the CS+E and CS+N were attenuated, whereas there was no such correlation for SCRs to the CS−. Following tryptophan depletion, individuals more intolerant of uncertainty showed significantly less emotional expression to all three stimuli during the spontaneous recovery phase. Importantly, the SCR to the US was unaffected by ATD. ATD also attenuated responses during the reinstatement and reacquisition phases, consistent with the spontaneous recovery phase results, however there was no longer evidence of differential conditioning.

## Discussion

The aim of this study was to advance the understanding of how serotonin influences the retention of conditioned emotional reactions. Here we showed, for the first time, that pharmacologically modulating serotonin affected the expression of aversive emotional memory in humans. During the key phase of the study - spontaneous recovery - depletion diminished arousal responses to the CS+s and CS- non-specifically and differential conditioning was preserved. Analysis of individual subject plasma samples, however, revealed that a greater degree of depletion was associated with reduced emotional responding to the CS+s, with no effect on CS- responses. These plasma level findings suggest that aversive emotional memory was attenuated. Examining self-reported intolerance of uncertainty, a trait measure that has previously been related to spontaneous recovery (Dunsmoor et al., 2015), was critical for uncovering how ATD affected emotion by contributing to the prediction of the general linear model. Individuals who reported being more intolerant of uncertainty at baseline, furthermore, showed even lower responses during spontaneous recovery when depleted. ATD also attenuated emotional responding during the reinstatement and reacquisition phases, consistent with the spontaneous recovery results. Importantly, responses to the US were unaffected by ATD, indicating that the effect was specific to learned cues and not a general blunting of arousal encompassing responses to aversion itself. Mood was unaffected by depletion, consistent with previous ATD studies of healthy volunteers (Ruhe et al. 2007). By using a task that elicited physiological reactions, however, it was possible to uncover an effect of serotonin on emotion.

The primary implication of the study is that serotonin transmission is critical for conditioned threat memory expression. This signal may be boosted in individuals highly intolerant of uncertainty, and excessive serotonin signalling may be an important contributor to the persistence of pathological emotional reactions. We reinforce and extend the observation that intolerance of uncertainty contributes to spontaneous recovery (Dunsmoor et al., 2015): by implicating serotonin signalling in this phenomenon, the present data suggest that trait intolerance of uncertainty may be a latent marker of vulnerability to serotonergic dysregulation.

Surprisingly few studies have investigated the influence of serotonin on threat conditioning processes in humans (Bauer, 2015). The effects of dietary/pharmacological manipulations aimed at lowering serotonin have been studied primarily in relation to the acquisition of threat conditioning in humans (Hensman et al., 1991; Hindi Attar et al., 2012; Robinson et al., 2012). The current study represents an important extension of this work by addressing a critical clinically relevant question: how does lowering serotonin impact the intensity with which previously formed emotional memories return? Indeed, aberrant spontaneous recovery, assessed using experimental paradigms analogous to the present one, has been demonstrated most notably in PTSD (Milad et al., 2009), as well as in OCD (McLaughlin et al., 2015; Milad et al., 2013) and schizophrenia (Holt et al., 2012). These disorders are commonly treated with drugs acting on serotonin (Stahl, 2013). This study may therefore inform both pathophysiology and mechanisms underlying treatment.

The directionality of the depletion effects - a reduction rather than enhancement of emotion - may seem counterintuitive. These results, however, are in line with and advance influential theories of serotonin function (Cools et al., 2011; Deakin, 2013; Deakin & Graeff, 1991), and are consistent with an array of experimental data (Bauer 2015; Bocchio et al., 2016; Deakin, 2013; Hensman et al., 1991; Hindi Attar et al., 2012). Serotonin is thought to be critically involved in predicting punishment, and aversively conditioned cues stimulate serotonin release (Bauer 2015; Deakin, 2013; Deakin & Graeff, 1991; Bocchio et al., 2016). The present results are most directly comparable to, and therefore substantiated by, two studies on the role of serotonin in healthy volunteers that also employed SCR to measure threat conditioning of neutral cues. Hensman et al. (1991) showed that the 5-HT2C and −2A receptor antagonist ritanserin - which diminishes the effects of serotonin signalling - impaired conditioning. Likewise, Hindi Attar et al. (2012) used ATD to show the same pattern: an attenuation of conditioning. These autonomic responses closely paralleled functional magnetic resonance imaging (fMRI) data indicating that ATD diminished signals in the amygdala and orbitofrontal cortex that were otherwise evoked by cues predictive of aversion (Hindi Attar et al., 2012). Our observation of diminished threat responses in the spontaneous recovery phase, in conjunction with the findings from Hensman et al. (1991) and Hindi Attar et al. (2012), therefore strengthens the punishment prediction framework of serotonin function advanced by Deakin & Graeff (1991).

The results of this study appear to agree with what is known about the basic serotonergic innervation of different amygdala subnuclei. The basolateral nucleus of the amygdala (BLA) is critical for storing associations between cues and aversive outcomes (LeDoux, 2000). The central nucleus of the amygdala (CeA), meanwhile, is a major source of output from the amygdala and signals downstream to structures including the hypothalamus and periacqueductal grey that contribute to defensive reactions such as perspiration in humans and freezing in rodents (LeDoux, 2000). Critically, the BLA receives dense serotonergic innervation whilst the CeA receives weak serotonergic input (Bauer 2015). This is remarkably consistent with the present findings: emotional responses to predictive cues (CSs), which should heavily engage serotonin signalling in the BLA, were modulated by ATD, whereas SCR to the aversive outcome itself (shock; US) was unaffected. Indeed, it is the CeA (which receives weak serotonergic input) that responds to aversive outcomes (Michely et al., 2019). That activity associated with aversive expectations occurs in the BLA, but not the CeA, has furthermore been associated with individual differences in trait anxiety in humans (Michely et al., 2019).

One limitation of the study is that complete extinction on Day 1 was not achieved. An experimental design with two CS+s, only one of which was to be extinguished, was used with the goal of comparing retention of conditioning alone (the CS+N; not extinguished) versus the retention of extinction (the CS+E; extinguished), two different processes that ultimately could not be definitively parsed. Whilst the lack of differential response between the CS+E and CS+N on Day 2 is likely due to incomplete extinction on Day 1, it is also possible, based on Bouton (2002), that the inclusion of the CS+N (not extinguished) in all phases of Day 2 cued memory for conditioning on Day 1 - thus enhancing memory expression for CS+E - more so than an extinction memory trace. On Day 2, SCR habituated in the phases after the critical test of spontaneous recovery, which often occurs (Gershman & Hartley 2015), but which made it more difficult to ascertain effects on reinstatement and reacquisition.

Another limitation of the study is that serotonin was not measured directly, and instead a common proxy measure involving plasma levels of tryptophan was used. Tryptophan is the amino acid precursor of serotonin and ATD has been shown to produce temporary reductions in central serotonin synthesis in humans (Nishizawa et al. 1997). Whilst the validity of ATD as a selective method to study serotonin has been questioned (van Donkelaar et al. 2011), this perspective has been rebutted on the basis of considerable evidence (Crockett et al. 2012a; Young 2013). The case that ATD reduces central serotonin is bolstered by consonant results from human studies employing ATD and rodent experiments that induce profound serotonin loss using the neurotoxin 5,7-DHT. “Waiting impulsivity” is a prime example, which can be induced in humans following ATD (Worbe et al. 2014), and in rats after serotonin depletion via 5,7-DHT (Winstanley et al. 2004).

The present study contributes to a surprisingly small literature examining how serotonin affects human threat conditioning (Bauer, 2015). We have shown for the first time that lowering serotonin attenuates the subsequent return of threat responses that were conditioned prior to depletion. This is a particularly important question from a clinical standpoint, and efforts to determine factors underpinning the retention of pathological emotional memories have been otherwise widespread (Graham and Milad, 2011; Milad and Quirk, 2012). We furthermore demonstrated - extending Dunsmoor et al. (2015) - that individual differences in self-reported intolerance of uncertainty were critical for understanding how serotonin affected spontaneous recovery. Greater intolerance of uncertainty was correlated with even lower spontaneous recovery following depletion. The use of such trait markers may be a valuable tool for refining which individuals may be most affected by serotonergic challenges. Integrating traits and neurochemical state is relevant for understanding vulnerability in healthy individuals, and may inform transdiagnostic mechanisms in clinical populations to refine psychiatric classification (e.g. Research Domain Criteria; Cuthbert & Insel, 2013) and help direct pharmacological strategies.

## Acknowledgements

This research was funded by a Wellcome Trust Senior Investigator Award (104631/Z/14/Z) awarded to T.W.R. B.J.S. receives funding from the National Institute for Health Research (NIHR) Cambridge Biomedical Research Centre (Mental Health Theme); the views expressed are those of the authors and not necessarily those of the NIHR or the Department of Health and Social Care. R.N.C.’s research is supported by the UK Medical Research Council (MC_PC_17213). J.W.K. is supported by a Gates Cambridge Scholarship. We would like to thank the staff at the NIHR/Wellcome Trust Clinical Research Facility at Addenbrooke’s Hospital, where the study was conducted, and Rachel Kyd of the Cambridge University Hospital Research & Development Office for assistance with study approval.

## Supplemental Material

### Inclusion criteria

Conditioning data from Day 1 were evaluated to determine whether participants qualified to continue with Day 2 of the conditioning experiment. Participants were said to have conditioned if they showed greater responses to both CS+s than to the CS−, averaged across either the entire acquisition phase, the first half of acquisition or the latter half of acquisition.

### Exclusion criteria

Exclusion criteria were summarized in the Methods. Psychiatric screening was conducted using the Mini-International Neuropsychiatric Interview (MINI; Sheehan, et al., 1998). Other exclusion criteria included pregnancy, past use of endocrine medication, use of St John’s Wort, regular consumption of over 38 units of alcohol per week, consumption of more than five cigarettes per day, use of cannabis more than once per month, use of other recreational drugs besides cannabis more than five times in the lifespan, cardiac or circulation problems, respiratory issues including asthma, gastrointestinal disorders, kidney disorders, thyroid problems, head injury, a bleeding disorder, and diabetes.

### Skin conductance

Electrodes were attached to the distal phalanges of the index and middle finger on the other arm to the shock electrode, which was counterbalanced. Base to peak increase in SCR to the CSs was valid if an increase began within a window of .5 to 4.5 seconds after stimulus onset. SCR to the CSs and US were low-pass filtered, smoothed, and square root transformed to normalize the distribution. SCR to CSs across all phases were divided by the average SCR to the US on Day 1 to enable between-subjects comparisons (e.g. Hartley et al. 2012).

### Statistics

Data were analysed using MATLAB (MathWorks) and SPSS (IBM). The Greenhouse-Geisser correction was used where applicable, in designs with within-subjects factors, to correct for violation of the sphericity assumption as determined by Mauchly’s test.

### Additional rating data

We collected rating data from 44 participants (n = 25 on depletion) on how much they enjoyed the task each day. Using a repeated measures ANOVA with serotonin status (placebo, ATD) and day (day 1, day 2) as factors, there was no main effect of ATD (F_(1,42)_ = 1.339, p = .254) nor was there a serotonin-by-day interaction (F_(1,42)_ = 0.492, p = .487). We also collected rating data from 37 participants (n = 20 on depletion) asking how uncomfortable they found the shocks on each day. Repeated measures ANOVA with group assignment (placebo versus ATD) and day (day 1, day 2) as factors showed there was no main effect of ATD (F_(1,35)_ = 0.148, p = .703) nor was there a group-by-day interaction (F_(1,35)_ = 0.339, p = .564).

